# Longitudinal neurochemical profiling after experimental stroke using edited MR spectroscopy

**DOI:** 10.64898/2026.09.10.750586

**Authors:** Ioana F. Grigoras, William T. Clarke, Mohamed Tachrount, Lauren Gill, Clémence Ligneul, Antoine Cherix, Myrto Lavda, Kamila Szulc-Lerch, Jason P. Lerch, Charlotte J. Stagg, Yvonne Couch

## Abstract

**Background:** Alterations in excitatory and inhibitory neurotransmission contribute to stroke pathology and recovery, yet longitudinal assessment of these changes *in vivo* remains limited. We investigated the feasibility of longitudinal edited proton magnetic resonance spectroscopy (^1^H-MRS) to quantify γ-aminobutyric acid (GABA) and glutamate following experimental focal cerebral ischaemia.

**Methods:** Male rats underwent endothelin-1-induced striatal ischaemia or sham surgery. Longitudinal edited ^1^H-MRS was performed at baseline and 2-, 7- and 30-days post-stroke at 7-tesla with a MEGA-sLASER sequence. A voxel was positioned within the ipsilesional motor cortex adjacent to the lesion. Behavioural outcomes (grip strength, adhesive removal and CatWalk gait analysis) were assessed alongside spectroscopy, and immunohistochemistry for Iba1, GFAP and ICAM-1 was performed at 30 days.

**Results:** Stroke induced a transient reduction in GABA+/tCr within the ipsilesional motor cortex, with significantly lower concentrations than sham animals at 7 days (p<0.05), returning to baseline by 30 days. Glutamate/tCr and GABA/glutamate ratio were unchanged throughout the study. Behavioural deficits were mild, with significant effects on motor outcomes such as grip strength, gait swing speed and paw placement, while sensory areas of the brain seemed largely unaffected. Histological analysis revealed no persistent differences in cortical Iba1, GFAP or ICAM-1 immunoreactivity at 30 days.

**Conclusions:** Longitudinal ^1^H-MRS is a feasible approach for monitoring dynamic changes in inhibitory neurochemistry following experimental stroke. The transient reduction in peri-lesional GABA, despite minimal behavioural impairment and an absence of persistent cortical inflammation, supports the use of edited spectroscopy as a translational tool to investigate neurochemical mechanisms underlying network dynamics in post-stroke recovery.

## Introduction

Ischemic stroke induces complex alterations in excitatory and inhibitory neurotransmission that contribute to acute injury and post-stroke recovery. Glutamate-mediated excitotoxicity is a well-established feature of early ischemic injury, while alterations in γ-aminobutyric acid (GABA) signalling may influence cortical plasticity and functional recovery during later stages of repair.^1^ Because GABAergic signalling is thought to regulate the capacity of surviving neural circuits to reorganise after injury, longitudinal assessment of GABA may provide insight into mechanisms of recovery that are not captured by structural imaging alone.^2, 3^

Proton magnetic resonance spectroscopy (^1^H-MRS) non-invasively measures brain metabolites to provide a tool for monitoring neurochemical alterations after stroke,^4, 5^ including low-concentration metabolites such as GABA. Although clinical and pre-clinical studies have reported altered glutamatergic and GABAergic signalling following stroke, longitudinal characterisation across acute and subacute phases remain limited, as do studies specifically looking at GABA in pre-clinical models.^6, 7^

In the present study, we used longitudinal edited ^1^H-MRS^8^ to quantify GABA+ and glutamate following endothelin-1-induced ischaemia in rats. Metabolic changes were assessed alongside behavioural outcomes and histological measures to investigate the evolution of post-stroke neurochemical alterations over time.

## Materials and Methods

### For full methodological details see *Supplementary Data*

Briefly, male Wistar Han rats (Inotiv; 100–125 g on arrival) underwent surgery to induce focal ischaemia by stereotaxic injection of 1 µl of endothelin-1 (ET-1; 25 pmol/µl) into the left striatum. Sham-operated animals underwent identical procedures with vehicle (0.9% saline). All animals underwent behavioural testing and magnetic resonance scanning. Magnetic resonance imaging and edited ^1^H MR spectroscopy of the left motor cortex (Fig. 1F) were performed on a 7T Bruker scanner under isoflurane anaesthesia with continuous physiological monitoring. GABA+macromolecule (GABA+) concentrations were quantified from difference spectra, while glutamate was quantified from edited-off spectra. Metabolite levels were referenced to total creatine, tCr ([tCr] = [Cr] + [PCr]). At *in vivo* study completion, animals were culled and brain tissue was processed for histology. Immunohistochemistry was performed using antibodies against ionized calcium-binding adaptor molecule-1 (Iba-1), glial fibrillary acidic protein (GFAP), and intercellular adhesion molecule-1 (ICAM-1). Longitudinal neurochemical data were analysed using linear mixed-effects models in R (lme4, lmerTest, emmeans), with intervention group and timepoint as fixed effects, subject as a random effect, and baseline values included as covariates. Post-hoc comparisons were performed using the Kenward–Roger method.^9^ Behavioural and histological data were analysed using mixed-model analysis of variance (ANOVA) with Bonferroni-corrected post-hoc comparisons where appropriate using Graphpad Prism (v11.0.0). Statistical significance was defined as p<0.05.

**Figure 1.**
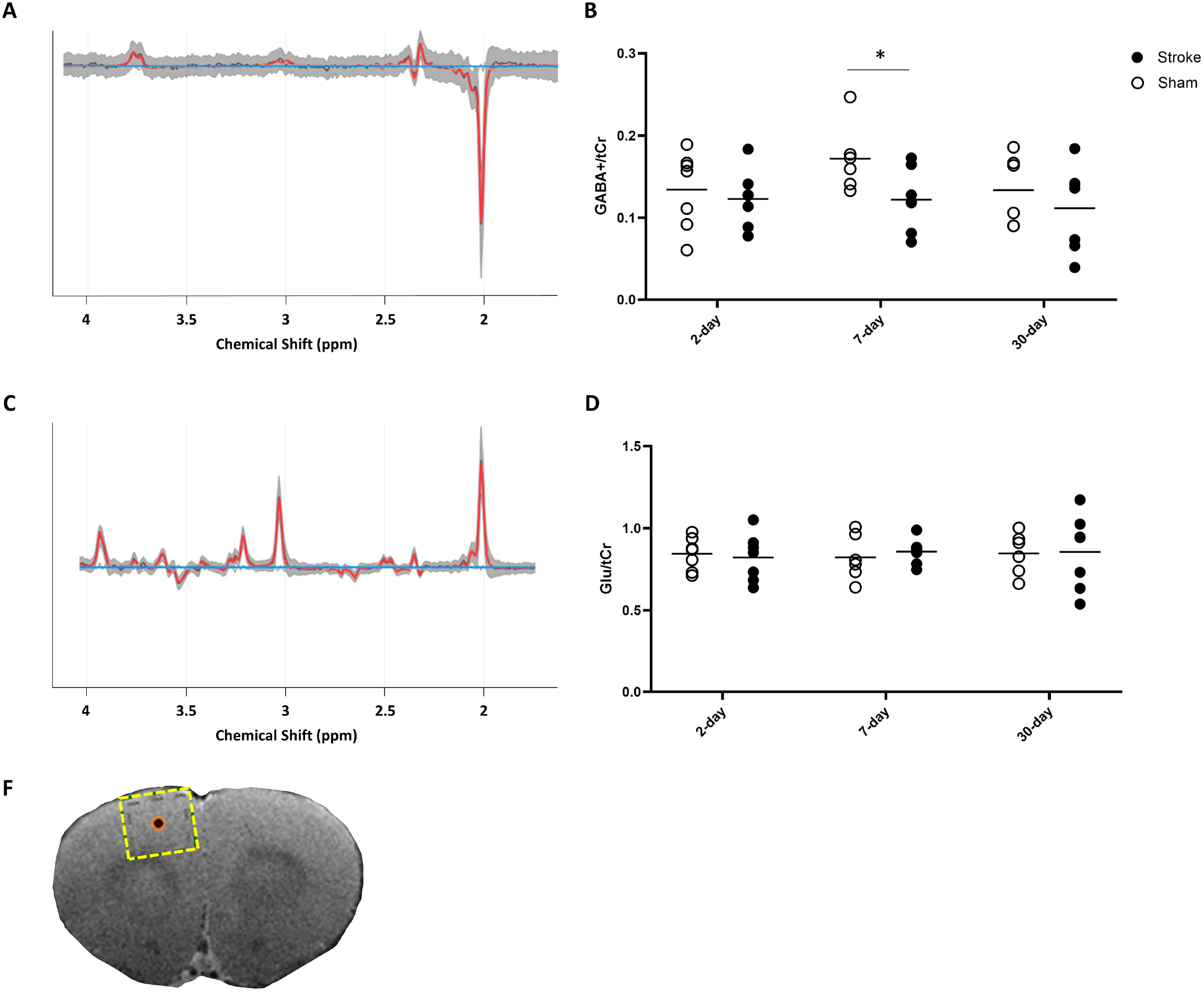
*Proton MR Spectroscopy*. (A and C) show mean spectra (black line, ± 1 standard deviation – shaded area), and mean model fits (red line), for the difference condition, targeting [GABA]), and the edit-OFF condition, targeting [glutamate]. (B and D) show [GABA] and [glutamate] concentrations (expressed as a ratio to [tCr]) for each subject at each time point post-intervention. (F) shows the voxel placement in a coronal section, for saggital and axial placement see Supplementary Information. Individual data points are shown and lines represent the mean. *p < 0.05.

## Data Sharing and Availability

The data supporting the findings of this study are available from the corresponding author upon reasonable request. Analysis code for the magnetic resonance spectroscopy data is available at https://git.fmrib.ox.ac.uk/grigoras/mrs_stroke_rodent

## Results

### MRS detects subtle changes in GABA, but not glutamate, after ischaemia

Longitudinal MRS analysis revealed a transient reduction in GABA+/tCr (Fig.1A & B) in a lesion-adjacent and functionally connected region of the brain following stroke. No main differences of time post-stroke or stroke vs sham were established in a mixed-effects model but post-hoc analysis demonstrated that stroke animals exhibited significantly lower GABA+/tCr ratios than sham animals at 7 days post-intervention (p<0.05; Fig. 1B). No significant differences were observed at 2 or 30 days. In contrast, glutamate/tCr (Fig.1C & D) did not differ significantly across time or between groups at any timepoint (Fig. 1D).

### Ischaemia in the striatum results in minimal motor impairment in the rat

Striatal ischaemia had a significant main effect on bilateral grip strength (mixed-effects ANOVA; p<0.05; Fig.2A) but there were no post-hoc differences between groups. There was no effect of ischaemia on sensation, as measured by the sticky tape test (Fig.2B). CatWalk gait analysis revealed subtle locomotor deficits following focal striatal stroke (Fig. 2C–F). Swing speed (the speed of the paw during the swing phase of the gait cycle, providing a measure of limb advancement during locomotion) showed a significant main effect of stroke (p<0.05; Fig. 2C), although post-hoc comparisons did not identify differences at individual time points. Stride length demonstrated a significant effect of time (p<0.05; Fig. 2D), with no effect of stroke or stroke × time interaction. Similarly, max contact area (the maximum paw area in contact with the walkway surface at any point during the stance phase, providing a measure of paw loading and weight-bearing during walking) trended towards being affected by stroke (p=0.052; Fig. 2E) but with no individual differences post-hoc. Mean contact intensity was unaffected by stroke or time (Fig. 2F). The changes in grip strength and the reduction in max contact area in the absence of changes in mean contact intensity suggests that striatal stroke altered fine motor control of paw placement without substantially affecting limb loading, consistent with a mild impairment of motor control rather than marked weakness.

**Figure 2.**
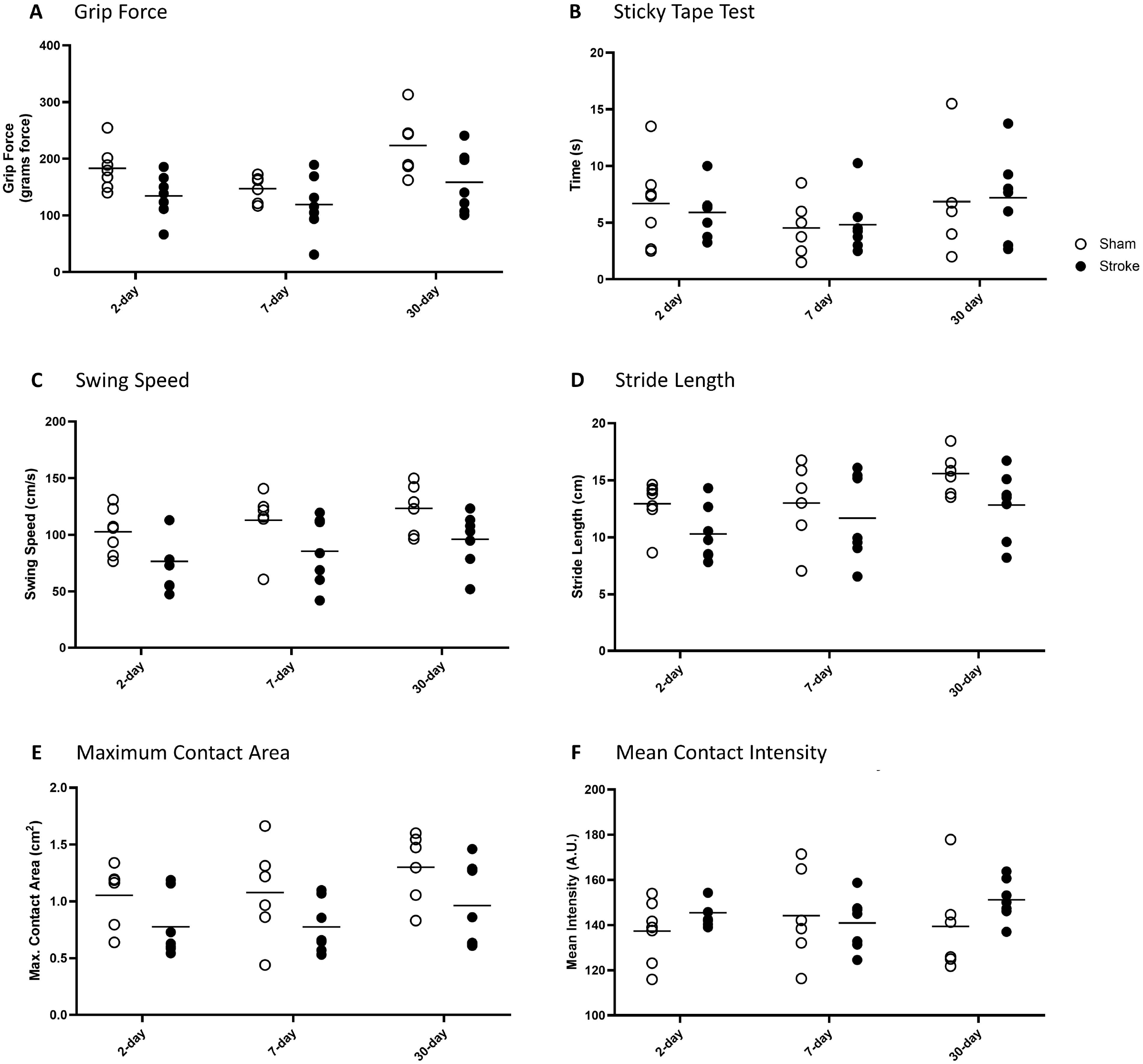
Longitudinal behavioural assessment following endothelin-1-induced striatal ischaemia. (A) Bilateral grip strength demonstrated a significant main effect of stroke. (B) Adhesive removal (sticky tape) test showed no significant differences between groups. CatWalk gait analysis of the contralateral forepaw assessed (C) swing speed, which showed a significant main effect of stroke; (D) stride length, which showed a significant effect of time only; (E) maximum contact area, which demonstrated significant effects of stroke, time and the stroke × time interaction, with a reduction in stroke animals at 7 days post-stroke; and (F) mean contact intensity, which was unchanged between groups. Individual data points are shown and lines represent the mean. *p < 0.05.

**Figure 3.**
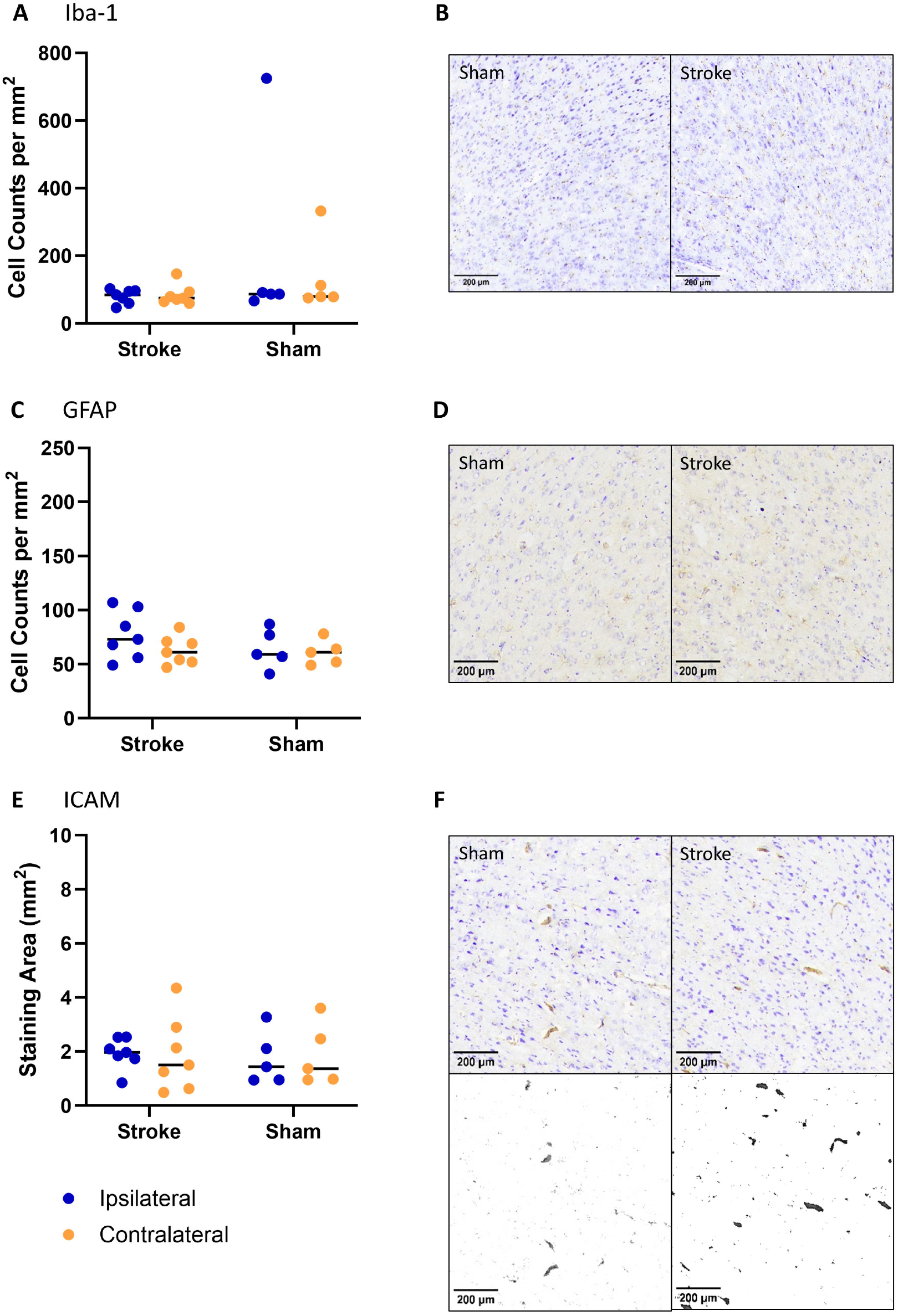
Histological assessment of neuroinflammation in the ipsilesional motor cortex 30 days following endothelin-1-induced striatal ischaemia. (A) Quantification of Iba1-positive microglia revealed no significant difference between stroke and sham animals. (B) Representative images of Iba1 immunoreactivity in the ipsilesional motor cortex of sham and stroke animals. (C) Quantification of GFAP-positive astrocytes demonstrated no significant difference between groups. (D) Representative images of GFAP immunoreactivity in the ipsilesional motor cortex of sham and stroke animals. (E) Quantification of ICAM-1 immunoreactivity, measured as percentage threshold-positive area, showed no significant difference between stroke and sham animals. (F) Representative images of ICAM-1 immunostaining in the ipsilesional motor cortex of sham and stroke animals. Individual data points are shown and lines represent the mean, scale bars represent 200 µm.

### Striatal ischaemia does not result in motor cortex inflammation at 30 days

Immunohistochemistry was carried out at 30 days, a time when the lesion had largely resolved. In the region of the motor cortex, where the MRS was carried out, there was no significant effect of stroke on Iba-1 cell counts, suggesting no change in microglial activity in this region (Fig.3A & B). Similarly, there were no effects of stroke or time on the number of GFAP-positive astrocytes in the motor cortex in stroke animals compared to shams (Fig.3C & D). ICAM expression was measured by threshold area staining and was not found to be different at 30 days post-stroke when compared to shams (Fig.3E & F).

## Discussion

To our knowledge, this is the first study demonstrating *in vivo* spectroscopy to measure glutamate as well as GABA in a region of the brain not directly affected by the stroke lesion. Our preliminary data demonstrate that endothelin-1 causes mild motor deficits and that MR spectroscopy can reveal changes in GABA and glutamate in the motor cortex, a region functionally connected to the area of the stroke.

While proton MRS has been widely used in pre-clinical stroke research, previous studies have predominantly focused on conventional metabolites, with comparatively little attention given to longitudinal assessment of inhibitory neurotransmission *in vivo*. Where neurotransmitters such as GABA and glutamate have been measured it has often been in an *ex vivo* setting,^6^ or it has been measurements of glutamate^7^ or other neurochemicals,^10, 11^ in the absence of GABA, which typically requires spectral editing for reliable detection *in vivo*. GABAergic inhibition plays a central role in regulating cortical excitability and synaptic plasticity.^2, 12, 13^ Indeed, we have shown that changes in inhibitory tone have been proposed to facilitate post-stroke remodelling and motor recovery.^2^ Longitudinal measurement of GABA may, therefore, provide insight into neurochemical processes underlying recovery that are not captured by conventional structural imaging. One limitation of this technique is the use of total creatine as an internal reference, as creatine metabolism may itself be altered following stroke. However, water referencing is also potentially confounded by post-stroke changes in tissue water content, and as such complementary referencing approaches should be considered in future studies.

In this study, GABA concentrations were reduced at 7 days following stroke but had returned to levels comparable with sham animals by 30 days, suggesting that the observed changes reflected a dynamic phase of post-stroke adaptation rather than persistent disruption of inhibitory neurotransmission. This reduction in GABA occurred during the same post-stroke period in which the most pronounced behavioural deficits were observed. Given the established role of GABAergic signalling in regulating cortical excitability and experience-dependent plasticity,^13^ these findings are consistent with the hypothesis that transient alterations in inhibitory tone accompany early functional reorganisation within surviving motor networks following subcortical injury.^2^ Future studies, beyond the scope of the current work, would benefit from specific-motor learning tasks carried out within this plastic period post-stroke, correlated with measurements of GABA and therapeutic interventions designed to restore excitation-inhibition balance.^12^

A strength of the approach used here is the flexibility of voxel placement to address different neurobiological questions. In the current study, spectroscopy was performed in the motor cortex overlying the striatal lesion, allowing longitudinal assessment of neurochemical changes within a cortical region involved in motor control and post-stroke plasticity. Given the central role of GABAergic signalling in regulating cortical excitability and motor learning, this approach provides insight into adaptive changes occurring within surviving motor networks following subcortical injury. Future studies could extend this methodology to alternative lesion–voxel combinations. For example, following focal motor cortical stroke, edited MRS could be used to investigate secondary neurochemical changes within the striatum, where the loss of corticostriatal glutamatergic input may alter the activity of the predominantly GABAergic neurons that form the principal output of the basal ganglia. Such studies would provide a means of investigating reorganisation post-stroke and demonstrate the utility of both the endothelin-1 model and the use of pre-clinical MR spectroscopy studies to determine network level changes after brain injury.

One limitation of our study is that our histological outcomes were, by necessity, taken at the final time-point but histological analysis at day 7 could provide additional information. Previous studies using the same model have shown that inflammatory responses evolve over the first week following stroke, before transitioning towards tissue remodelling and repair.^14^ Others have demonstrated GFAP immunoreactivity at 7 days in the regions bordering a striatal lesion.^15^ Importantly, the majority of studies measure immunoreactivity within the core, our data are in the perilesional area. Measurements within the ischemic lesion do show significant increases in astrocyte and vascular reactivity and have been included in the supplementary file (Supplementary Figure S4).

Reactive astrocytes play an important role in regulating neurotransmitter homeostasis through glutamate uptake, glutamate–glutamine cycling and metabolic support of neuronal function, and transient astrocyte activation during the subacute phase of stroke could therefore contribute to alterations in tissue neurotransmitter pools detected by MRS. Further studies, including those introducing drugs to manipulate GABA,^12^ would enable us to confirm or refute the cellular contributions to our spectroscopic signal.

Whilst the present study demonstrates the feasibility of combining longitudinal ^1^H-MRS with behaviour and histology in a pre-clinical model of focal ischaemia, it does have several additional limitations beyond the above mentioned histological endpoint. Only male animals were included, and future studies should determine whether similar neurochemical changes occur in females, given known sex differences in stroke pathophysiology and recovery.^16^ Similarly, age^17^ and comorbidities,^18, 19^ known contributors to stroke outcomes, were not taken into account

Pre-clinical studies have traditionally been used to search for lesion-reducing therapeutics, with little attention going on long-term recovery. This has generated a translational gap in stroke research.^20^ By studying rodent models longitudinally and using translational techniques, such as proton spectroscopy, which are also applicable in humans we can hope to bridge this gap.

## Conclusion

This study demonstrates the feasibility of longitudinal *in vivo* edited ^1^H-MRS to quantify GABA and glutamate following experimental focal cerebral ischaemia. These preliminary findings support the application of edited 1H MRS to investigate changes in excitatory–inhibitory balance during stroke progression and recovery and provide a foundation for future studies in larger cohorts.

## Acknowledgements

For the purpose of open access, the author has applied a CC BY public copyright licence to any Author Accepted Manuscript version arising from this submission.

## References

1. Clarkson AN, Huang BS, Macisaac SE, Mody I, Carmichael ST. Reducing excessive gaba-mediated tonic inhibition promotes functional recovery after stroke. Nature. 2010;468:305–309

2. Blicher JU, Near J, Naess-Schmidt E, Stagg CJ, Johansen-Berg H, Nielsen JF, et al. Gaba levels are decreased after stroke and gaba changes during rehabilitation correlate with motor improvement. Neurorehabil Neural Repair. 2015;29:278–286

3. Puts NA, Edden RA. In vivo magnetic resonance spectroscopy of gaba: A methodological review. Prog Nucl Magn Reson Spectrosc. 2012;60:29–41

4. Wang X, Li YH, Li MH, Lu J, Zhao JG, Sun XJ, et al. Glutamate level detection by magnetic resonance spectroscopy in patients with post-stroke depression. Eur Arch Psychiatry Clin Neurosci. 2012;262:33–38

5. Johnstone A, Levenstein JM, Hinson EL, Stagg CJ. Neurochemical changes underpinning the development of adjunct therapies in recovery after stroke: A role for gaba? J Cereb Blood Flow Metab. 2018;38:1564–1583

6. Baranovicova E, Kalenska D, Lehotsky J. Glutamate/gaba/glutamine ratios in intact and ischemia reperfusion challenged rat brain subregions, the effect of ischemic preconditioning. Metab Brain Dis. 2025;40:121

7. Yoo CH, Baek HM, Song KH, Woo DC, Choe BY. An in vivo proton magnetic resonance spectroscopy study with optimized echo-time technique for concurrent quantification and t2 measurement targeting glutamate in the rat brain. MAGMA. 2020;33:735–746

8. Mescher M, Merkle H, Kirsch J, Garwood M, Gruetter R. Simultaneous in vivo spectral editing and water suppression. NMR Biomed. 1998;11:266–272

9. Kenward MG, Roger JH. Small sample inference for fixed effects from restricted maximum likelihood. Biometrics. 1997;53:983–997

10. Jimenez-Xarrie E, Davila M, Gil-Perotin S, Jurado-Rodriguez A, Candiota AP, Delgado-Mederos R, et al. In vivo and ex vivo magnetic resonance spectroscopy of the infarct and the subventricular zone in experimental stroke. J Cereb Blood Flow Metab. 2015;35:828–834

11. Shemesh N, Rosenberg JT, Dumez JN, Muniz JA, Grant SC, Frydman L. Metabolic properties in stroked rats revealed by relaxation-enhanced magnetic resonance spectroscopy at ultrahigh fields. Nat Commun. 2014;5:4958

12. Grigoras IF, Geist E, Johnstone A, Clarke WT, Emir U, Nettekoven C, et al. Baclofen, a gabab receptor agonist, impairs motor learning in healthy people and changes inhibitory dynamics in motor areas. Imaging Neurosci (Camb). 2025;3

13. Stagg CJ, Bachtiar V, Johansen-Berg H. The role of gaba in human motor learning. Curr Biol. 2011;21:480–484

14. Abeysinghe HC, Bokhari L, Dusting GJ, Roulston CL. Brain remodelling following endothelin-1 induced stroke in conscious rats. PLoS One. 2014;9:e97007

15. Al-Khishman NU, Qi Q, Roseborough AD, Levit A, Allman BL, Anazodo UC, et al. Tspo pet detects acute neuroinflammation but not diffuse chronically activated mhcii microglia in the rat. EJNMMI Res. 2020;10:113

16. Roy-O′Reilly M, McCullough LD. Age and sex are critical factors in ischemic stroke pathology. Endocrinology. 2018;159:3120–3131

17. McCullough LD, Mirza MA, Xu Y, Bentivegna K, Steffens EB, Ritzel R, et al. Stroke sensitivity in the aged: Sex chromosome complement vs. Gonadal hormones. Aging (Albany NY). 2016;8:1432–1441

18. Manwani B, Liu F, Scranton V, Hammond MD, Sansing LH, McCullough LD. Differential effects of aging and sex on stroke induced inflammation across the lifespan. Exp Neurol. 2013;249:120–131

19. Lyden PD, Bosetti F, Diniz MA, Rogatko A, Koenig JI, Lamb J, et al. The stroke preclinical assessment network: Rationale, design, feasibility, and stage 1 results. Stroke. 2022;53:1802–1812

20. Neuhaus AA, Couch Y, Hadley G, Buchan AM. Neuroprotection in stroke: The importance of collaboration and reproducibility. Brain. 2017;140:2079–2092

