## Supplemental File for "Longitudinal neurochemical profiling after experimental stroke using edited MR spectroscopy"

**Supplementary Material**

*Methods*

*Animals*: Male Wistar Han rats (Inotiv; 100–125 g on arrival) were housed under standard laboratory conditions with *ad libitum* access to food and water and a 12-hour light/dark cycle. All procedures were approved by the UK Home Office and local Animal Welfare and Ethical Review Body (PPL PP7444704). Animals underwent baseline assessments 48 hours prior to intervention and follow-up at 2, 7, and 30 days post-intervention. One animal did not recover following the 2-day MRI scan and was excluded from subsequent longitudinal analyses.

*Surgery*: Focal ischemic stroke (n=7) was induced by stereotaxic injection of 1 µl of endothelin-1 (ET-1; 25 pmol/µl) into the left striatum under isoflurane anaesthesia (2% induction, 1.5% maintenance; 60/40 O_2_/N_2_O) which resulted in a focal sub-cortical lesion (Fig.S1). Sham-operated animals (n=7) underwent identical procedures with vehicle (0.9% saline). Surgeries were performed by an independent operator and all subsequent testing (imaging and behaviour) were performed blind.

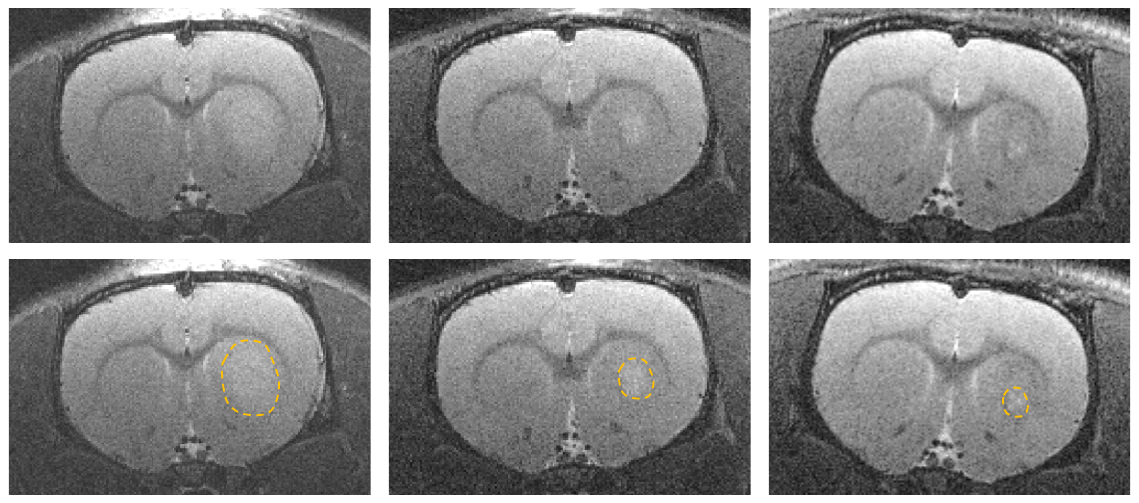

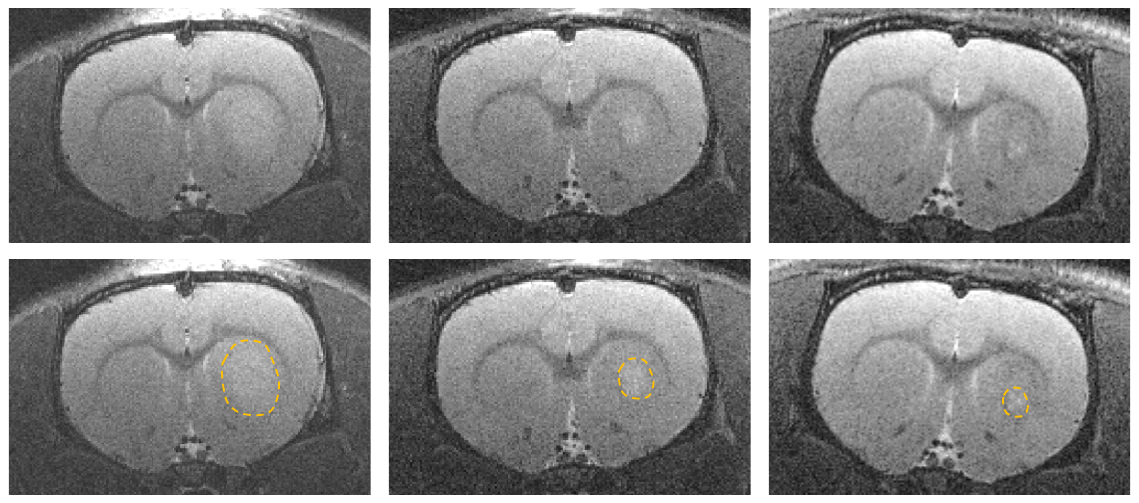

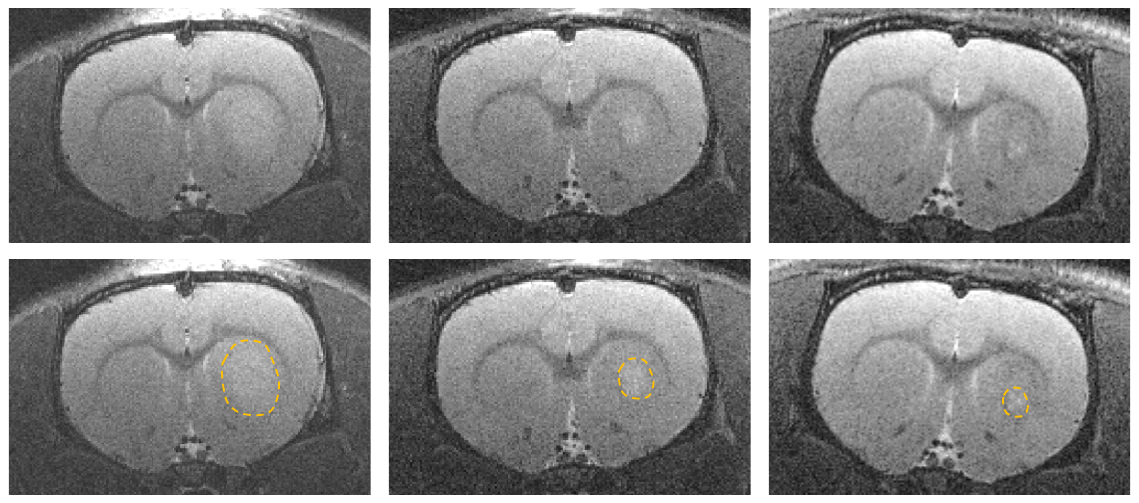

2 days

7 days

30 days

2 days

7 days

30 days

Figure S1. Representative structural MRI demonstrating the evolution of the endothelin-1-induced lesion over time. Top row: Representative T2-weighted anatomical MR images from a single stroke animal acquired at 2, 7 and 30 days following endothelin-1-induced striatal ischaemia. Hyperintense signal within the ipsilateral striatum is evident at 2 days and progressively decreases over time, consistent with resolution of the lesion. Bottom row: Same representative T2-weighted images with the lesion highlighted. Images are representative of the temporal evolution observed across the study cohort.

Magnetic Resonance Protocol:

During the MR scanning, rats were put under isoflurane anaesthesia with continuous physiological monitoring. The scans were performed on a 7T Bruker (Ettlingen, Germany) BioSpec 70/20 USR scanner using a 72mm 1H volume transmit coil and a 2x2 rat brain surface receive array coil (Bruker, Ettlingen, Germany). After adjusting the position of the rat inside the magnet based on fast localisation scans and acquiring the field map, axial, sagittal, and coronal high in-plane spatial resolution T2 weighted scans were acquired with the following acquisition parameters: TE/TR = 33/2500 ms, echo spacing = 11 ms, RARE factor = 8, spatial resolution = 80x80x600 um3, 2 averages and scan time of 2min40s each. Based on these structural images, the MRS voxel was manually positioned in the left motor cortex above the endothelin-1-induced striatal lesion using anatomical landmarks, with placement optimised to minimise overlap with white matter and the cortical boundary and to avoid the lesion (Figure S2). The high order shimming was performed within a prescribed volume that encompasses the voxel of interest. The edited 1H magnetic resonance spectroscopy data were acquired using a custom implementation of MEGA-sLASER sequence [2]. Full details of the MR spectroscopy protocol alongside example spectra are provided in the attached MRSinMRS (Supplementary Table 1), but briefly acquisition parameters were: voxel size 1.8 × 2.3 × 2.8 mm³; TE/TR = 68/3000 ms; editing pulse offset 1.9 ppm (on) and 7.5 ppm (off)). Three or four 20-minute acquisition blocks were collected per session alongside a water reference scan. Voxel placement was in the motor cortex above the lesion

Spectroscopy data were processed using FSL-MRS.^1^ Raw Bruker data were converted to NIfTI-MRS format,^2^ coil-combined, aligned, averaged, separated by editing dimension, prior to final alignment and subtraction to generate a difference spectrum . GABA+macromolecule (GABA+) concentrations were quantified from difference spectra, while glutamate was quantified from edit-off spectra. Metabolite levels are reported as a ratio to total creatine, tCr ([tCr] = [Cr] + [PCr]).

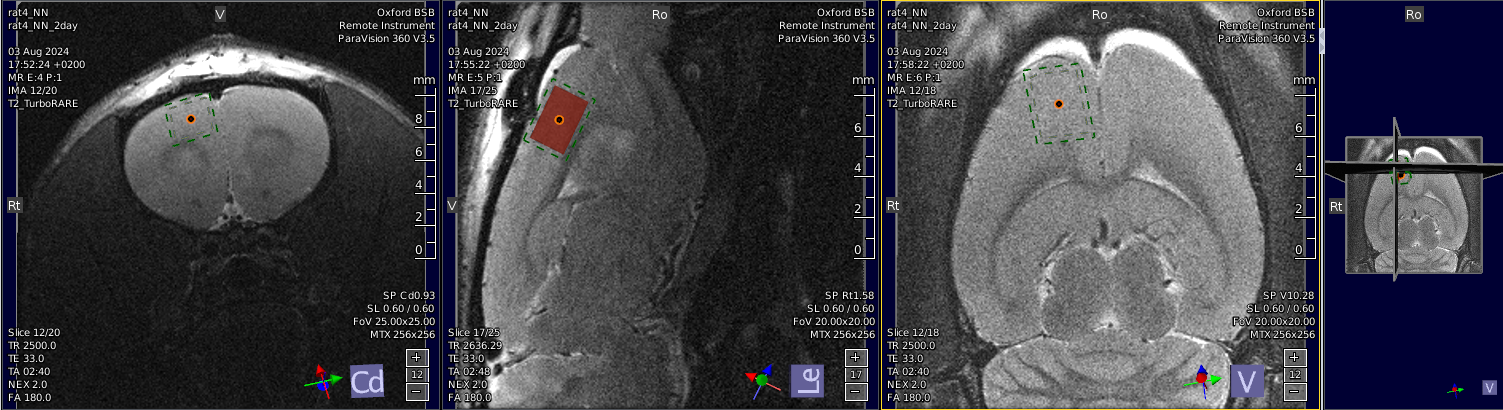

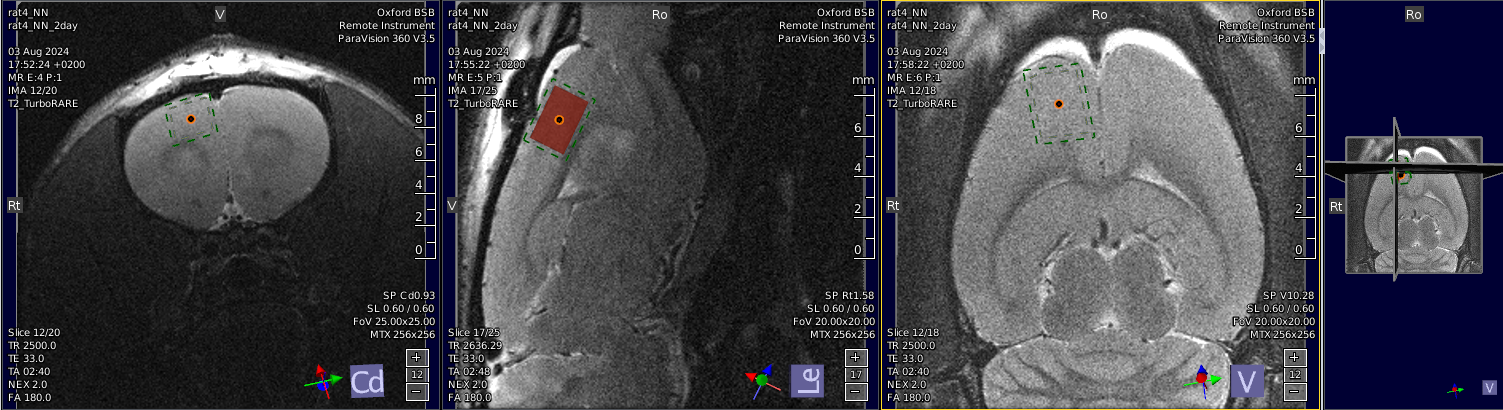

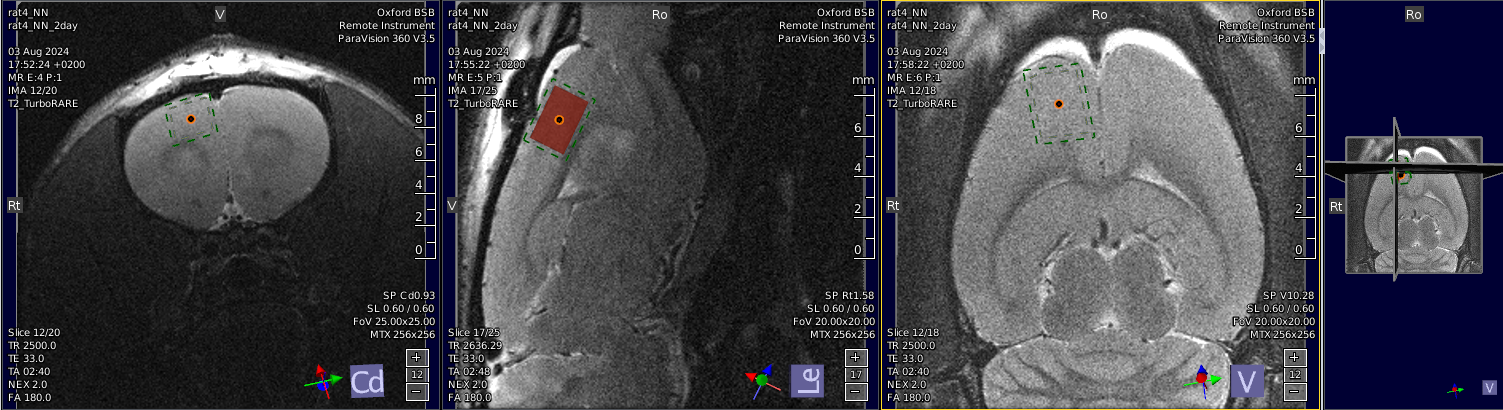

Figure S2. Representative voxel placement for ^1^H-MRS acquisition. Representative T2-weighted anatomical MR image showing the location of the spectroscopy voxel (1.8 × 2.3 × 2.8 mm³) positioned within the ipsilesional motor cortex adjacent to the endothelin-1-induced striatal lesion. Voxel placement was performed following structural imaging and was maintained consistently across animals and imaging sessions.

*Behaviour*: Behavioural testing was carried out at baseline, two days, seven days and thirty days post-ischemia. Grip strength testing was performed on both paws simultaneously using a grasping grid and the grip strength meter (Harvard Apparatus, UK). A total of three to five tests for each rat were performed per time point and an average taken for the five tests. Sticky tape testing was performed using a 1cm x 1cm piece of fabric band aid (Elastoplast) applied to the contralateral forepaw. Animals were placed in an empty cage and recorded on video until they had successfully removed the tape. Post-hoc the time to notice (touch) the affected paw and to remove the sticky tape were recorded. CatWalk gait analysis was performed using the CatWalk XT automated gait analysis system (Noldus Information Technology, Wageningen, The Netherlands). Rats were allowed to traverse the illuminated glass walkway voluntarily, and only uninterrupted runs meeting predefined speed and variation criteria were included for analysis. Three compliant runs were recorded for each animal and averaged. Gait parameters were calculated using CatWalk XT software, including run duration, average speed, regularity index (the percentage of normal step sequence patterns relative to the total number of paw placements, providing a measure of interlimb coordination during locomotion), and paw-specific measures of max contact area (the maximum paw area in contact with the walkway surface at any point during the stance phase), print area (the total area of the paw contacting the walkway during a step, providing a measure of paw contact and loading during locomotion), stride length (the distance between successive placements of the same paw, providing a measure of step progression during locomotion), duty cycle (the percentage of the step cycle during which the paw is in contact with the walkway, providing a measure of the relative duration of limb loading during locomotion), and mean intensity (the mean intensity of the reflected paw print during walkway contact, reflecting the degree of paw pressure/loading applied to the walkway). For unilateral stroke animals, analyses focused on the contralateral (right) forepaw and hindpaw. All data for Catwalk analysis, including ipsilateral paw and all other measures not included in the paper, have been included in this supplementary file.

*Tissue Processing*: At study completion, animals were euthanised by pentobarbital overdose followed by transcardial perfusion with phosphate-buffered saline and 4% paraformaldehyde (PFA). Brains were harvested, post-fixed in 4% PFA for 24 hours, and cryoprotected in 30% sucrose for min. 48 hours prior to embedding in optimal cutting temperature compound. Coronal brain sections (12 µm) were cut using a Leica CM3050 cryostat and mounted onto gelatin-coated slides for subsequent histological analyses.

*Histology*: Immunohistochemistry was performed on coronal brain sections using antibodies against ionized calcium-binding adaptor molecule-1 (Iba-1 1:500; AbCam; ab178847), glial fibrillary acidic protein (GFAP 1:500; AbCam ab7602, Cambridge, UK), and intercellular adhesion molecule-1 (ICAM-1 1:1000; Sigma-Aldrich/eBioscience 12-0549-42). Sections were processed using a streptavidin–biotin peroxidase method with Vectastain ABC kits (Vector Laboratories, UK) according to the manufacturer’s instructions. Immunoreactivity was visualised using 3,3′-diaminobenzidine (DAB) chromogen, followed by cresyl violet counterstaining. Stained sections were imaged using an EVOS light microscope. Image analysis was performed using ImageJ (National Institutes of Health, Bethesda, MD, USA) within the region of the motor cortex used for imaging, and within the lesion area (Fig.S3). For Iba-1 and GFAP, cell number was counted within a pre-defined area. For ICAM staining, images were acquired under identical microscope settings and analysed using a consistent threshold applied to all sections within the experiment. Following thresholding to identify positive immunostaining, the total area of ICAM-positive staining within the region of interest was measured and expressed as the percentage area of positive staining. Area-based quantification was selected in preference to cell counting because ICAM immunoreactivity is associated with vascular structures of variable size and morphology, as well as non-vascular cellular elements following cerebral ischaemia and as such counting is a less appropriate quantification technique.

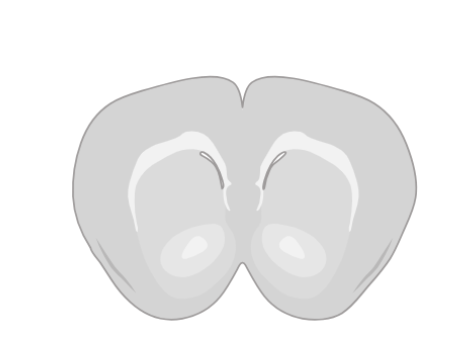

Figure S3. Outline of regions analysed for histology.

*Statistics:* Longitudinal neurochemical and behavioural data were analysed using linear mixed-effects models in R (lme4, lmerTest, emmeans), with intervention group and timepoint as fixed effects, subject as a random effect, and baseline values included as covariates. Post-hoc comparisons were performed using the Kenward–Roger method. Histological data were analysed using analysis of variance (mixed-effects model ANOVA) with Bonferroni-corrected post-hoc comparisons where appropriate. Statistical significance was defined as p<0.05.

*Results*

*Striatal ischaemia results in persistent local inflammation at 30 days*

Quantification of immunohistochemistry was also performed in the core lesion area. Similar to the motor cortex, there was no significant effect of stroke on Iba1-positive microglial cell counts within the striatum (Fig. S4A & B). However, stroke induced a significant increase in the number of GFAP-positive astrocytes (p<0.05; Fig. S4C & D), indicating persistent astrocytic activation within the lesion core. ICAM-1 expression, quantified by threshold area staining, demonstrated significant main effects of stroke (p<0.05) and brain region (ipsilateral versus contralateral; p<0.01), with Bonferroni post hoc analysis revealing significantly increased ICAM-1 immunoreactivity in the ipsilateral striatum of stroke animals compared with sham controls (p<0.05; Fig. S4E & F).

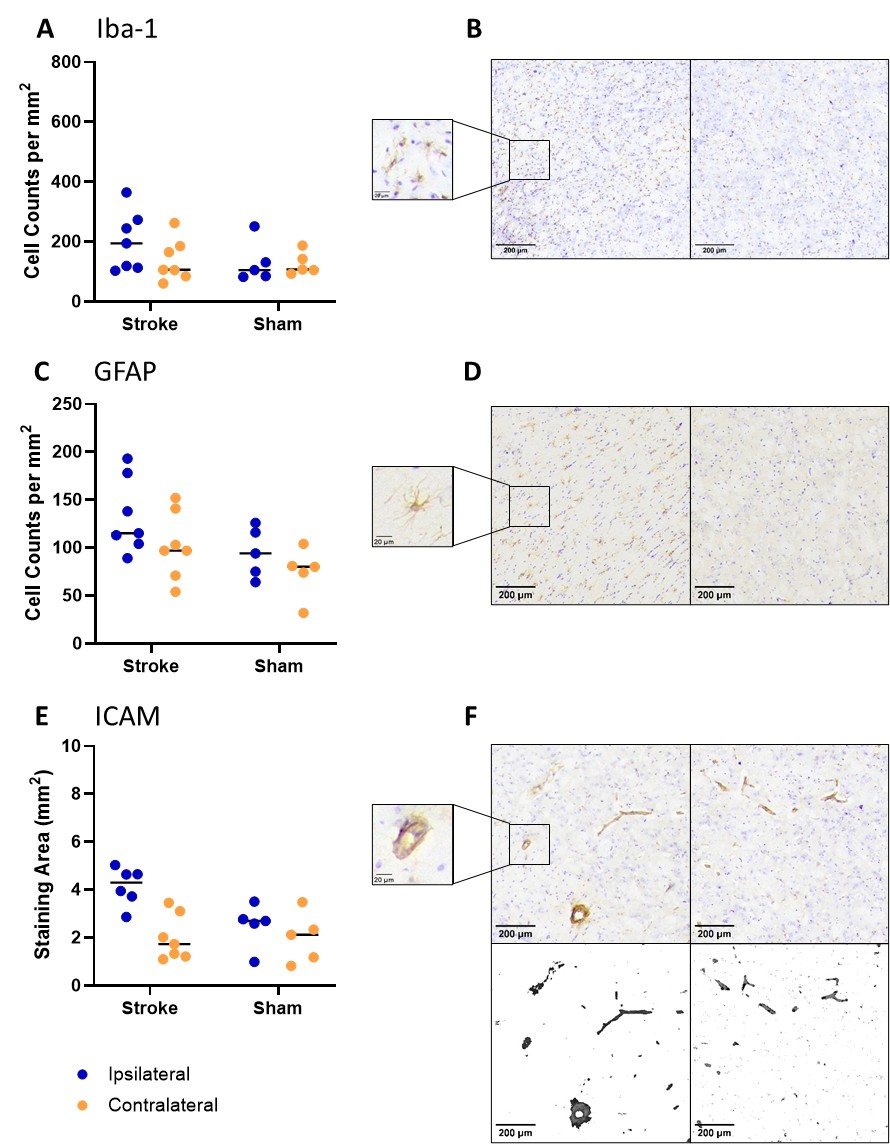

Figure S4*. Histological assessment of neuroinflammation in the ipsilesional striatum 30 days following endothelin-1-induced striatal ischaemia.* (A) Quantification of Iba1-positive microglia revealed no significant difference between stroke and sham animals. (B) Representative images of Iba1 immunoreactivity in the ipsilesional motor cortex of sham and stroke animals. (C) Quantification of GFAP-positive astrocytes demonstrated no significant difference between groups. (D) Representative images of GFAP immunoreactivity in the ipsilesional motor cortex of sham and stroke animals. (E) Quantification of ICAM-1 immunoreactivity, measured as percentage threshold-positive area, showed no significant difference between stroke and sham animals. (F) Representative images of ICAM-1 immunostaining in the ipsilesional motor cortex of sham and stroke animals. Individual data points are shown and lines represent the mean, scale bars represent 200 µm.

*References*

1.	Clarke WT, Stagg CJ, Jbabdi S. Fsl-mrs: An end-to-end spectroscopy analysis package. *Magn Reson Med*. 2021;85:2950-2964

2.	Clarke WT, Bell TK, Emir UE, Mikkelsen M, Oeltzschner G, Shamaei A, et al. Nifti-mrs: A standard data format for magnetic resonance spectroscopy. *Magn Reson Med*. 2022;88:2358-2370
